# Computational identification of splicing phenotypes from single cell transcriptomic experiments

**DOI:** 10.1101/2020.11.04.368019

**Authors:** Yuanhua Huang, Guido Sanguinetti

## Abstract

RNA splicing is an important driver of heterogeneity in single cells, both through the expression of alternative transcripts and as a major determinant of transcriptional kinetics. However, the intrinsic coverage limitations of scRNA-seq technologies make it challenging to associate specific splicing events to cell-level phenotypes. Here, we present BRIE2, a scalable computational method that resolves these issues by regressing single-cell transcriptomic data against cell-level features. We show that BRIE2 effectively identifies differential alternative splicing events that are associated with a disease. Additionally, BRIE2 allows a principled selection of genes (differential momentum genes) that capture heterogeneity in transcriptional kinetics and improve quantitatively RNA velocity analyses. BRIE2, therefore, extends the scope of single-cell transcriptomic experiments towards the identification of splicing phenotypes associated with biological changes at the single-cell level.

## Background

Single-cell RNA-sequencing (scRNA-seq) has rapidly become the key technology to disentangle transcriptional heterogeneity in cell populations. Over the last five years, scRNA-seq has been successfully applied both to identify discrete cell states or subpopulations in normal or diseased tissues, e.g. [1, 2], and to infer continuous stages in cellular processes, e.g., pseudo-time [3] and cell differentiation [4]. More recently, scRNA-seq has further been applied to multi-sample designs with different donors, tissues, diseases or treatments. These experiments enable the discovery of cell type specific marker genes [5] or key pathways that are associated with the meta labels [2].

Beyond gene-level information, RNA processing within a gene also holds rich information for both categorical cell states and continuous cell differentiation. A key RNA processing step is splicing, where a precursor mRNA (pre-mRNA or un-spliced RNA) is spliced by removing intronic, non-coding regions, resulting in mature mRNA (or spliced RNA). Alternative splicing of exons further extends the molecular feature space, greatly contributing to cellular heterogeneity. A variety of studies have found that the abundance of splicing isoforms enables the identification of cell states [6] or disease conditions [7]. Additionally, the intrinsic kinetics of splicing provide a footprint of cellular dynamics during cell differentiation, which has motivated the recent flourishing of RNA velocity studies [8, 9] and time-series scRNA-seq on metabolically labelled nascent RNAs [10, 11, 12, 13].

Despite the fundamental role of RNA splicing, stochasticity in splicing is much less understood than that of gene-level expression, primarily due to the technical difficulties in recovering splicing information from scRNA-seq data. First, scRNA-seq data is highly sparse, particularly for droplet-based protocols including the popular 10x Genomics platform. This high sparsity, along with minimal initial molecule counts, leads to very high technical noise in scRNA-seq data, e.g., seemingly mono-isoform pattern [14], hence requiring careful statistical modelling. Second, splicing adds new layers of complexity to scRNA-seq analyses, and the requirements to quantify relative abundances of isoforms from indirect observations of fragment counts creates considerable computational difficulties. For all these reasons, the level of heterogeneity in splicing between different cells has been difficult to quantify. Perhaps more importantly, the identification of single-cell level splicing phenotypes, including alternative splicing events associated with a disease or genetic changes and genes with variable unspliced ratios across cell population, has been largely unfeasible, hindering an understanding of the role of splicing changes and aberrations in cellular state.

In this work, we study ratio of spliced vs unspliced RNAs and that of two alternative splicing isoforms (e.g., exon inclusion and exclusion) in a unified way, interchangeably termed as splicing ratio. Here, we address these above computational issues by directly incorporating the association of splicing phenotypes within the splicing quantification task itself. We introduce BRIE2, a Bayesian hierarchical model that predicts the splicing ratio from a set of features associated with cell-type/ state, as well as with the specific splicing event to be quantified. This enables us to robustly identify genes or splicing events that are associated with each cell level feature, while controlling and quantifying in a Bayesian manner the uncertainty from the noise and sparsity of the data. We show on simulated and real data sets that BRIE2 yields better quantification of splicing ratios and more effective detection of differential splicing between groups of cells, compared to state-of-the-art competitors. Additionally, by treating unspliced intronic as a form of alternative splicing, BRIE2 allows us to quantify differential transcriptional kinetics between cell types, thus providing us with an efficient way to select biologically relevant features for RNA-velocity analyses. We show on a number of examples that this procedure leads to more consistent and interpretable visualisations of biological process dynamics.

## Results and discussion

### Model Description

An unavoidable difficulty in splicing quantification from short-read protocols derives from the fundamental ambiguity of the data, as the vast majority of reads cannot be unambiguously assigned to a single isoform. This problem is compounded in scRNA-seq by the generally low number of reads, which frequently results in no unambiguous reads being mapped to a specific isoform. Therefore, using bulk-based methods or directly computing ratios of read counts assigned to specific isoforms is unlikely to effectively quantify the percentage of spliced-in (PSI) quantity for the majority of events in a given cell^[1]^. Our earlier work, BRIE [15] (from now on BRIE1), resolved this issue by regressing (suitably transformed) PSI values on sequence features through a Bayesian regression approach, therefore using genomic sequences to regularise and inform splicing predictions. This enables BRIE1 to transfer information across genes, identifying sequence features that are highly predictive of splicing efficacy in a particular cell, and providing a principled trade-off between imputation and data-driven estimation. However, because sequence features are normally the same between individual cells, BRIE1 is not particularly well suited to quantify differential splicing across cell types, and needs generally to be run independently on different cells, which can result in a significant computational burden.

BRIE2 starts again from a latent regression framework, but innovates over BRIE1 in two important ways: first of all, it augments the set of regressor features to include cell-specific features such as cell-type/ developmental stage (Fig. 1, Supplementary Fig. S1, and Methods). This enables us to statistically associate cell-level features with PSI-values associated with specific splicing events, thus defining quantitatively splicing as a single-cell level intermediate phenotype, but it considerably increases the complexity of the model (as data from all cells needs to be analyzed jointly). Second, to cope with the added complexity, BRIE2 is formulated as a variational discriminative model, thus enabling the use of advanced software (Tensorflow) and hardware (GPUs), and leading to orders of magnitudes in computational acceleration (>1,000 speed-ups; see Supplementary Fig. S2 and Methods).

**Figure 1.**
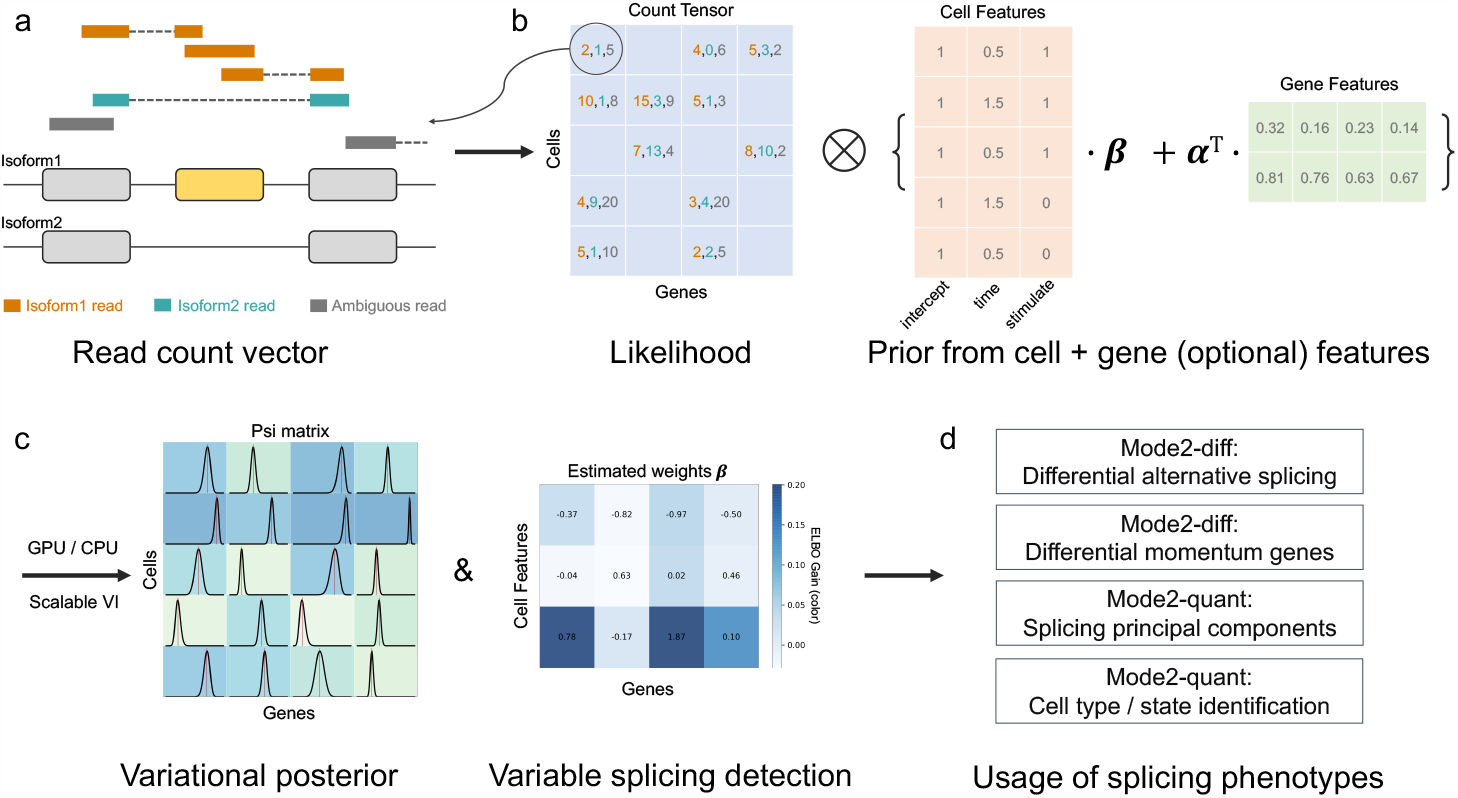
Illustration of BRIE2. (a) Reads are counted into isoform 1, isoform 2 or ambiguous groups according to its alignment identity, which constitutes a cell-by-gene-by-3 tensor. (b) The posterior distribution of isoform proportion PSI is defined by combining the likelihood from read counts and informative prior predicted by cell-level covariates and/or gene sequence features. (c) A `logit`-`normal` variational posterior and coefficients on covariates are optimised to approximate the exact posterior, where the evidence lower bound gain (ELBO) between including and excluding a certain cell feature set can be leveraged to select splicing phenotypes. (d) The selected differential splicing events or differential momentum genes on RNA velocity can be used as markers for downstream analysis, and the estimated PSI can be used for dimension reduction to enhance cell type/ state identification.

To recapitulate BRIE2’s operational capabilities, we report here the three major modes in which it can operate (see Table 1):

**Table 1.** The different usage modes of BRIE2.

|  | Features | Purpose |
| --- | --- | --- |
| Mode0 | None | Pure quantification with posterior |
| Mode1 | Gene features (genomic sequences, etc) | Imputation-enabled quantification |
| Mode2-quant | Unity cell features (i.e., intercept) | Quantification with aggregated prior |
| Mode2-diff | Cell features (condition, pseudo-time, etc) | Differential splicing detection |

Mode 0 in pure quantification mode, BRIE2 can simply perform quantification of exon inclusion ratios based on the available scRNA-seq reads, without using any auxiliary features. In this modality, BRIE2 is closely related to classical PSI quantification methods for bulk RNA-seq such as MISO [16], but with a different prior distribution (logit-normal centred on 0.5);

Mode 1 BRIE2 provides a much faster implementation of BRIE1, using sequence-based features associated with the event to regularise the estimation of PSI values and impute missing data;

Mode 2 BRIE2 can regress (logit transformed) PSI values against cell-level features, enabling the association of splicing phenotypes with cell types or with continuously varying cell features such as developmental time. Two specific examples of this mode are of particular relevance. The first one is to have a constant (unity) cell feature: in this case, the event-specific effect term will provide an adaptive prior over PSI which is informed by the splicing levels of the specific event across all cells. This mode (termed Mode 2-quant) is useful for regularised estimates of PSI values in a homogeneous cell population. Instead, differential quantification is obtained when using an indicator variable of cell type as a cell level feature: in this modality (Mode 2-diff) the event-specific effect term quantifies the effect of cell type on splicing variability for the specific event.

Multiple modes can be enabled at the same time, e.g. using simultaneously gene-level and cell-level features (combining modes 1 and 2-quant).

### Benchmarking BRIE2 on simulated data

BRIE1 was comprehensively benchmarked for its accuracy in identifying splicing ratios [15] against a variety of methods including Census [17], Cufflinks [18] and Kallisto [19]. BRIE2 also offers excellent quantification capabilities (see Supp. Fig. S4 for a comparison of BRIE1 with BRIE2 Mode 1 on estimating splicing percentages on real data). The quality of PSI estimates is considerably enhanced by the use of a unity feature (Mode 2-quant), as opposed to simple quantification (Mode 0), particularly for low coverage levels (see Supp. Fig. S5).

BRIE2’s main conceptual innovation is the availability of Mode 2-diff to associate splicing events with cell-level features. BRIE2 detects genes with differential PSI values by performing Bayesian model selection (see Methods) using the ELBO gain as a surrogate for Bayes factors. To assess its performance, we compare BRIE2 to two well-used methods for differential splicing detection in bulk RNA-seq, rMATS [20] and MAJIQ [21]. BRIE2 returns excellent performance in both sensitivity and specificity; Fig 2 shows precision-recall (PR) curves at three signal-to-noise ratios (magnitude of the effects) (see Supp. Fig. S6 for analogous ROC curves). At all levels, BRIE2 reports significantly better performance than either of the two competitors, demonstrating the efficacy of BRIE2 as a tool for detecting splicing changes across different cell types^[2]^.

**Figure 2.**
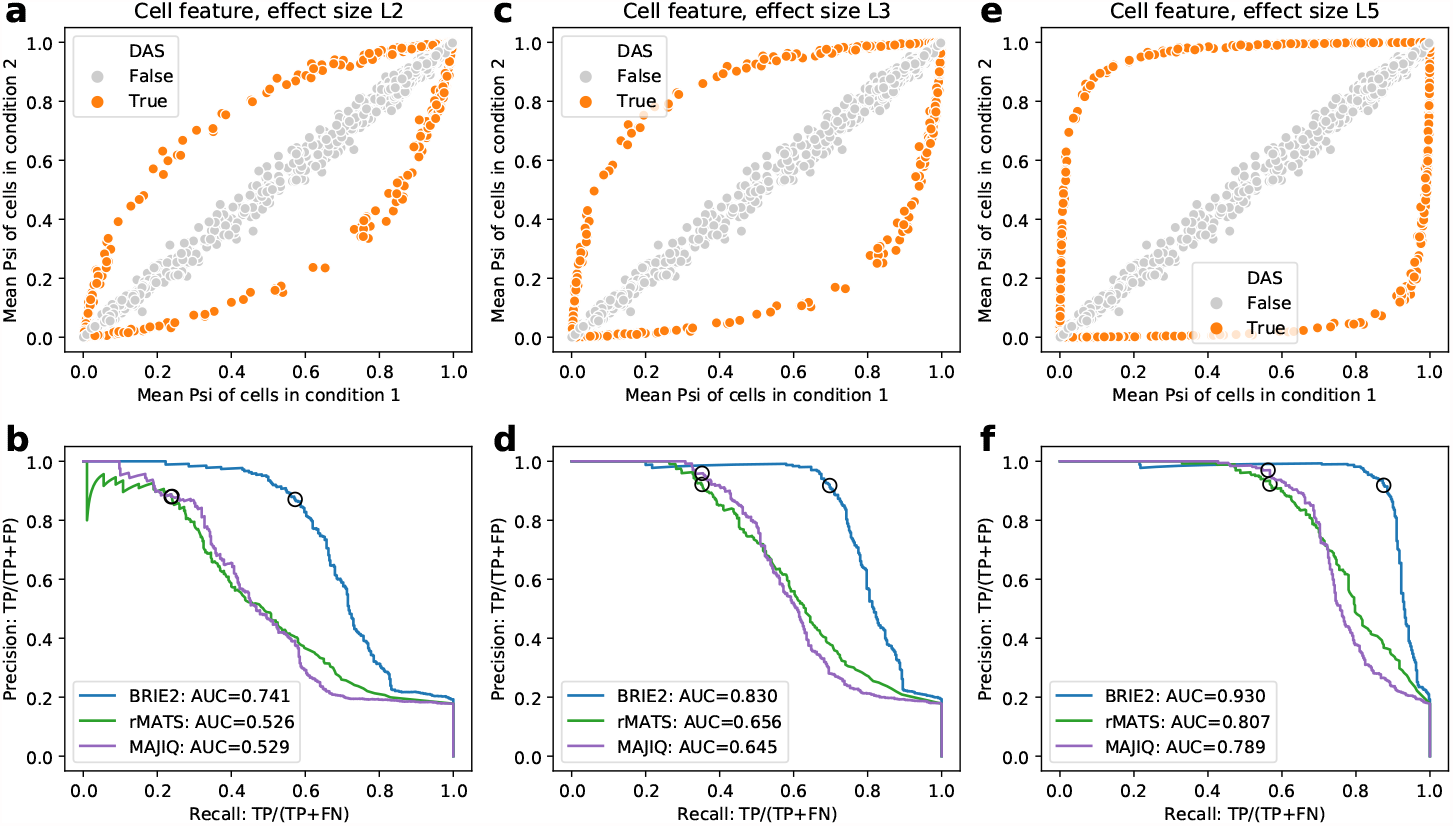
Evaluation of BRIE2 on detection of differential alternative splicing with simulated data. (a) Mean PSI of cells in condition 1 (x-axis) and condition 2 (y-axis), where 400 out of for 2,248 events are differential alternative splicing (DAS, coloured in orange). The effect size level L2 means effect size at logit scale is 2. (b) Precision-recall curve for detecting the DAS from simulated single-cell RNA-seq reads by BRIE2, rMATS and MAJIQ. The effect size is L2, as shown in (a). Black circles on the curve denote cutoff of `ELBO_gain` > 3 fo BRIE2 or |`delta_PSI`|> 0.2 for rMATS and MAJIQ. Similar to (a-b), more simulations and results are shown with effect size level L3 (c-d) and L5 (e-f), respectively.

### BRIE2 discovers hundreds of differential splicing events associated with multiple sclerosis

Next, we applied BRIE2 to analyse alternative splicing in multiple sclerosis, a neurological autoimmune disease. Falcão *et al* have generated 2,208 mouse cells using the SMART-seq2 protocol, with an equal number of cases (Experimental Autoimmune Encephalomyelitis, EAE mice) and controls [7]. Here, we analysed 3780 exon-skipping events that satisfied the quality control, e.g., more than 30 cells with unique reads, across 1,876 cells that have more than 3,000 total reads on the above events (Methods).

We first applied BRIE2 to quantify `PSI` using Mode 2-quant in Table 1. Note that the cell type information is not included for this initial PSI quantification. We collected this information in a `PSI` matrix (with dimensions number of cells times number of events), and performed a principal component analysis on it. We found that the `PSI` top principal components have strong cell-type specificity (see Fig. 3a-b for the first PC). This cell-type specificity of `PSI` PCs does not appear to be confounded by changes in gene expression (see Supp. Fig. S8).

**Figure 3.**
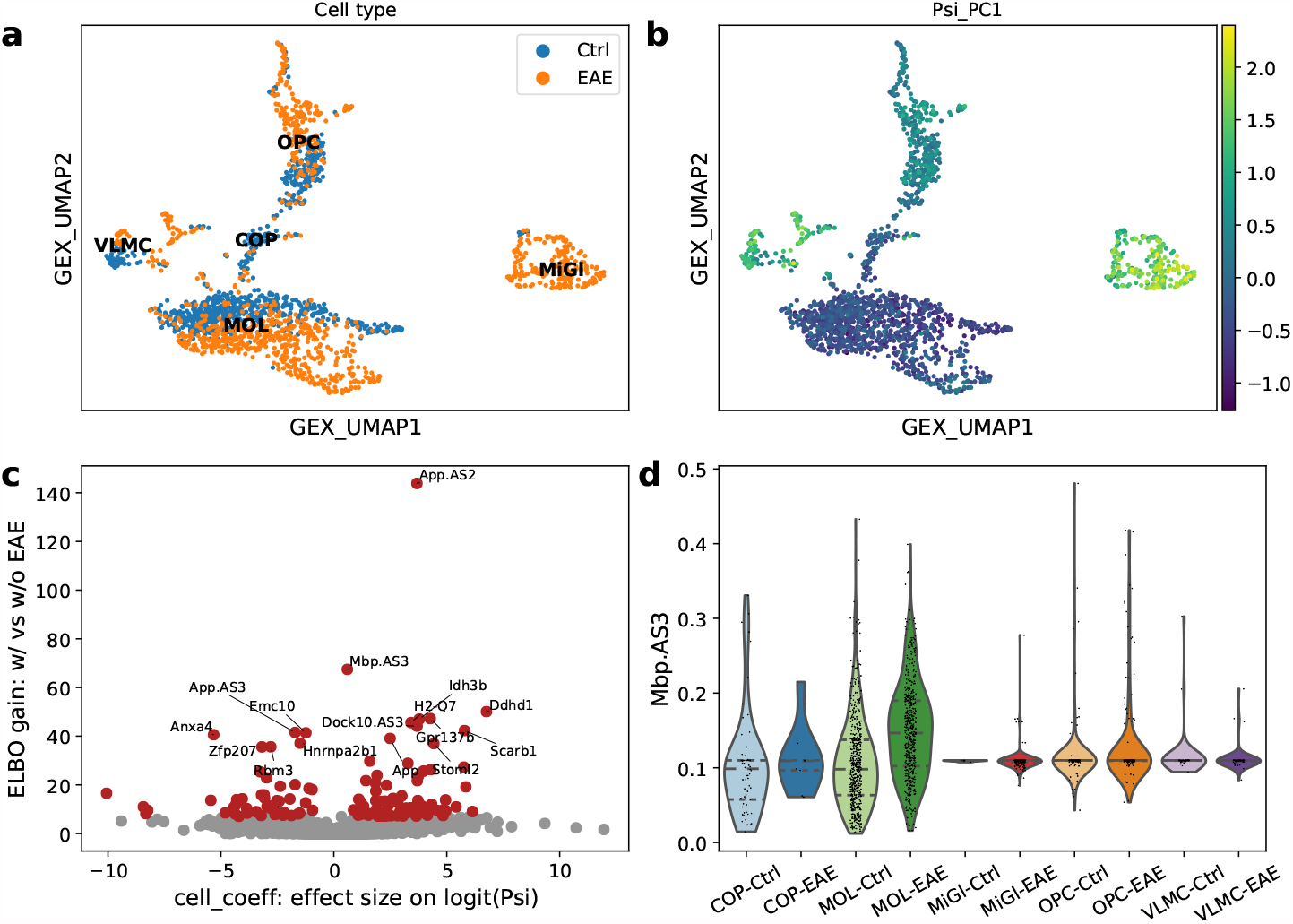
Differential splicing events on multiple sclerosis. (a) UMAP visualization of gene-level expression, annotated with cell types and EAE state. (b) UMAP coloured by the first principal component based on PSI matrix, which suggests that PSI PC has a global impact on cell types. (c) Volcano plot between -log10(FDR) and effect size on logit(PSI) for detecting differential splicing between EAE and control cells by BRIE2. (d) Violin plot on example gene Mbp (the exon3) for estimated PSI between EAE and control in each cell type. PSI values in panels (b) and (d) are quantified by only using a unity cell feature for aggregation, but not the EAE state label. EAE: Experimental Autoimmune Encephalomyelitis.

Additionally, we found that the top 20 `PSI` PCs can accurately predict the cell type (Supplementary Fig. S9a), and are reasonably predictive of disease state (AUC = 0.76, Supplementary Fig. S9b) on the most numerous cell type MOL. While using gene expression yields a considerably better prediction of disease state (AUC=0.95, Supp. Fig. S9b), using jointly splicing and expression covariates leads to a further (slight) improvement (AUC=0.96, Supp. Fig. S9b), indicating that some additional independent information is conveyed by splicing variables.

Running BRIE2 in Mode 2-diff, using disease state as a cell-level feature, detects 352 differential splicing events across 335 genes with ELBO_gain>4 that are associated with disease condition (Fig. 3c, Supplementary Fig. S10-11). Particularly, the myelin genes Mbp (ELBO_gain=67.4; Fig. 3d) and Pdgfa (ELBO_gain=12.8) are both identified as differential splicing events, which was highlighted in the original study [7] by using BRIE1. These events often have relatively few unambiguous reads and complex distributions of `PSI` values (see Supp. Fig. S10 and S11), including bimodality and long-tailed distributions. This level of variability in cell-level `PSI` values would make it difficult to adopt bulk-based strategies for differential splicing quantification (for example, by pseudo-bulking cells according to disease state).

### Differential momentum genes improve RNA velocity analyses

Global RNA-processing efficiency has recently been used to define the concept of RNA velocity associated with an individual cell [8, 9], which is rapidly becoming a major tool to study the dynamics of cellular processes at the single-cell level. The concept of RNA velocity is based on quantifying splicing kinetics by comparing the number of reads coming from pre-mRNA (unspliced) and mRNA, associating to each gene in each cell an RNA-processing speed which is then combined (and projected using any visualisation tool) to quantitate the dynamics of cellular processes at the molecular level. Implicitly, this is equivalent to treating spliced and unspliced RNAs as two different RNA conditions (equivalent as isoforms here).

Standard RNA velocity analyses are fully unsupervised, thus discarding available annotations during the (frequently crucial) step of selecting genes for velocity estimates. Instead, we propose to use BRIE2 to detect genes that have differential splicing ratios (spliced vs unspliced) associated with cell-level covariates, thus providing a biologically informed approach for selecting features to compute RNA velocities that are associated with cell transitions. We term these genes as *differential momentum genes* (DMG), as the differential splicing ratio implies a departure from the equilibrium between splicing and degradation rates (i.e., steady-state), likely due to changes in synthesis rate associated with changes in cell type. Therefore, the DMGs reflect the differential transcriptional activities between cell groups or states.

To see the impact of using DMGs in RNA-velocity analyses, we re-analyzed the data set of mouse Dentate Gyrus neurogenesis, introduced in [9], which well illustrates the impact of gene selection on cell transition inference. We used BRIE2 in Mode 2-diff to detect cell type-specific DMGs by using each cell type as the testing covariate and accounted for differences in coverage between cells by using gene detection rate as an additional cell-level covariate. We, therefore, examine the effect of using BRIE2 as a pre-selection step in velocity analyses, applying the same down-stream modelling to DMGs and default genes selected by the package scVelo [9]. The stochastic model is used here for illustrating that the differentiation direction can be corrected by using informative genes.

In Fig. 4a-b, we compare the cell differentiation paths inferred from RNA velocity based on the 634 genes selected by the package scVelo [9] and the 297 DMGs selected by BRIE2 (ELBO_gain>5 in any cell type), both selected out of the initial 3,000 quality-pass genes. While the overall picture is broadly in agreement, DMGs obtained from BRIE2 (Fig. 4b) highlighted a more obvious direction from oligodendrocyte precursor cells (OPCs) to myelinating oligodendrocytes (OLs) compared to scVelo either with its stochastic model (Fig. 4a) or dynamical model (Fig. 2 in [9]). Interestingly, this trajectory is not improved by using the differential kinetic gene sets detected by scVelo nor the differentially expressed genes detected edgeR (Supp. Fig. S12), while BRIE2’s performance is robust to the threshold choice of ELBO_gain (Supp. Fig. S13; read counts for top genes in Supp. Fig. S14). This observation remains even if zooming into the subset with only 103 OPC and OL cells (Supp. Fig. S15 and S16). We dwell here on the qualitative aspects of this comparison and their biological interpretation, but a fully quantitative assessment is provided in terms of cross-boundary correctness in Supp. Table S1, showing a marked improvement in using DMGs compared to other selection criteria.

**Figure 4.**
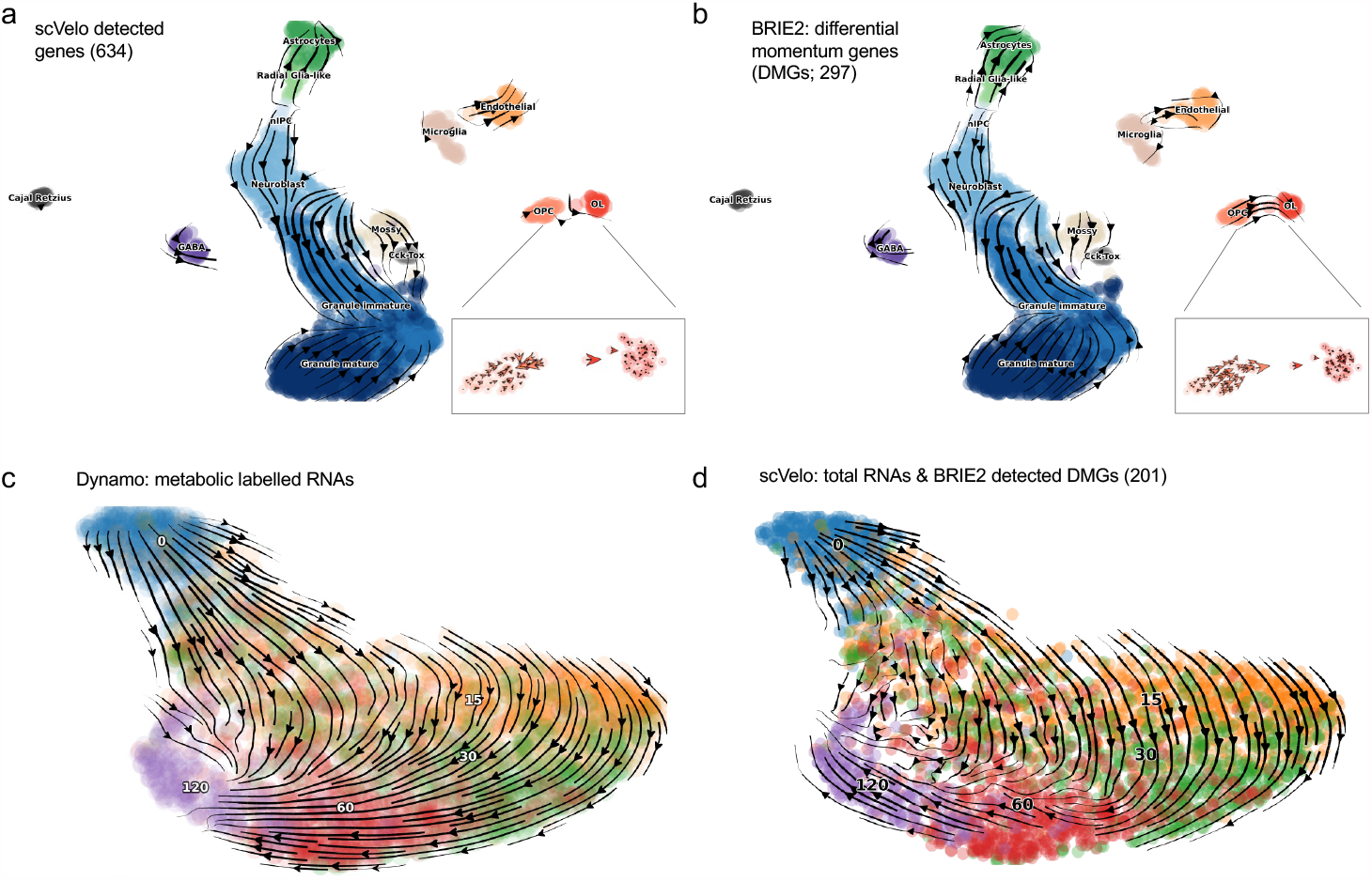
Differential momentum genes for RNA velocity. (a-b) Cell differentiation in neurogenesis inferred from RNA velocity by scVelo with different gene sets: (a) scVelo detected gene set requiring positive correlation between unlicensed and spliced RNAs; and (b) BRIE2 detected gene set that have differential spliced ratio in one cell type vs all others, which are termed as differential momentum genes. (c-d) State transitions of excitatory neurons inferred from RNA velocity with different methods: (c) Dynamo using metabolic labelling information measured by scNT-seq. (d) scVelo using total RNAs on 201 differential momentum genes detected by BRIE2. The colour denotes the time since stimulations: 0 (blue), 15 (orange), 30 (green), 60 (red) and 120 (purple) minutes.

Furthermore, we examined how the selection of DMGs improve the inference of cell state transition in time-series of neuronal scRNA-seq data generated by scNT-seq [13]. scNT-seq is a recently proposed technique where nascent RNAs are metabolically labelled, effectively providing a measurement of the age of a transcript. Using the information of metabolic labelling provides an effective ground truth and enables a consistent visualisation where cell transitions are strongly aligned with the time direction [22] (Fig. 4c). In the original paper, it was observed that such transitions are difficult to obtain only using the total RNAs; our own experimentation confirms that scVelo struggles to identify the right direction in the early stage of stimulation (i.e., 0 to 15 or 30min; Supp. Fig. S17-S18 for different settings; Supp. Table S1). Applying BRIE2 to detect DMGs by using the stimulation time as testing covariate, we found 421 DMGs significantly associated with time (ELBO_gain>5; Supp. Fig. S19-S20), with 201 genes overlapped with the top 2,000 highly variable genes selected by scVelo. By projecting the RNA velocity on these 201 DMGs, the cell transitions are largely corrected to the expected direction along the time (Fig. 4d; Supp. Table S1). This pattern remains even if varying the cut-off at ELBO_gain>7 for more stringent or ELBO_gain>3 for or more lenient DMGs (Supp. Fig. S21).

Taken together, these observations highlight the importance of feature selection when visualising cell transitions: in this light, DMGs detected by BRIE2 are likely to return more biologically informative angles, thanks to its detection of genes with differential transcriptional kinetics via the use of annotations.

## Conclusion

Splicing is a fundamental step in gene expression in higher eukaryotes and has the potential to represent an important intermediate phenotype in single-cell experiments. BRIE2 provides an effective and computationally efficient approach to link such intermediate phenotypes to cell-level covariates. Our results showed that BRIE2 identifies hundreds of splicing events linked to multiple sclerosis, and that inclusion of splicing events leads to improved cell-type classification on this translationally relevant data set.

While quantification of splicing events is certainly biologically important, it is likely to only be possible using technologies that sample evenly the transcriptome. Recent years, instead, have seen the increasing popularity of technologies that can upscale the number of cells assayed by sequencing only parts of the transcriptome (typically, the regions immediately preceding the polyA tail). Despite this enrichment, many such data sets still present a substantial number of intronic reads (presumably due to the abundance of repetitive A sequences within introns) which can be used to measure changes in RNA kinetics (so-called RNA velocity) and therefore provide a more accurate description of transitional cell states in large data sets. Our results showed that, in the presence of cell annotations, BRIE2 can be a useful tool to select relevant genes (differential momentum genes) which provide a smoother and more interpretable description of cell transitions within RNA velocity studies. The importance of selecting trajectory-informed genes for RNA velocity is also evidenced in another recent study [23].

Finally, while we believe BRIE2 to be a useful addition to the scRNA-seq analysis toolkit, it does have its limitations. The first one is intrinsic to the technology: splicing quantification can only, in general, be achieved with whole transcriptome sequencing technologies, ruling out popular approaches based on unique molecule identifier (UMI) and 3’ enrichment via polyA trapping. Secondly, BRIE2 focuses only on exon-skipping events, ruling out more complex architectural changes, or de-novo discovery of splicing variants. This is in contrast with many bulk methods, which are designed specifically to capture such complex events. While in principle application of similar ideas to scRNA-seq could lead to methods to detect such complex/ de-novo events even in single cells, we suspect that such methods will only become effective when single-cell technologies will start providing less sparse data.

## Methods

### Modelling of splicing isoform abundance

In this study, we jointly analyse *N* splicing events across *M* cells, focusing on two specific splicing events, for example, exon-skipping (SE) or genes with spliced vs unspliced RNAs (for the RNA-velocity part). For a splicing event *g* in a cell *c*, we use *ψ*_*c,g*_ to denote the fraction of a certain isoform; for conventional reasons, it refers to the isoform with exon-inclusion in SE event. Without loss of generality, we define the BRIE2 model on SE event here but it is applicable to any other two-isoform event.

In order to scale up the analysis across a large number of cells, reads aligned to a splicing gene are not modelled individually but rather aggregated into three groups depending on their isoform identity:

- group1: reads from isoform1 explicitly, e.g., on the junction between exon1 and exon2;
- group2: reads from isoform2 explicitly, e.g., on the junction between exon1 and exon3;
- group3: reads with ambiguous identity e.g., within exon3.

Thus, from the aligned reads file we could extract the count vector ***s***_*c,g*_ = [*s*_*c,g*,1_, *s*_*c,g*,2_, *s*_*c,g*,3_] for these three groups, with *n*_*c,g*_ = Σ_*k*∈1,2,3_ *s*_*c,g,k*_ as the total count. In addition, for each splicing event *g* we can pre-define the effective length *l*_*g,h,k*_, i.e., the (effective or corrected) number of positions in isoform *h* that can generate read being located in the region of read group *k*. This gene-specific 2-by-3 length matrix *L*_*g*_ can be defined from the exon structures encoded in the gene annotation, and the read counts are proportional to the effective lengths.

Given the total read counts *n*_*c,g*_ and its according effective length matrix *L*_*g*_, we can computed the likelihood of *ψ*_*c,g*_ (or equivalently its transformation *z*_*c,g*_ := `logit`(*ψ*_*c,g*_)) for observing the three-group reads counts ***s***_*c,g*_ as a multinomial distribution, whose proportion vector ***ρ***_*c,g*_ is coded by *ψ*_*c,g*_ and the effective length matrix *L*_*g*_ as follows,

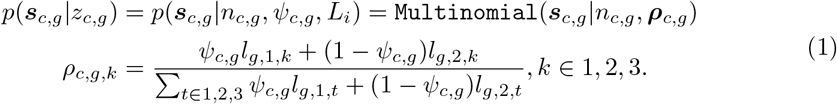

By definition, we have *l*_*g*,1,2_ = *l*_*g*,2,1_ = 0 and *l*_*g*,1,3_ = *l*_*g*,2,3_ for any splicing event *g*. Assuming conditional independence, we obtain the joint likelihood for all *N* splicing events in *M* cells by taking their product as follows,

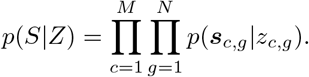

### Bayesian regression on splicing

In the BRIE2 model (see graphical representation in Supplementary Fig. S1), we aim to identify the regulatory factors of splicing from both gene level features ***x*** (e.g., splice site motif) and / or cell level features ***y*** (e.g., cell type) via a generalised linear model. Specifically, we assume that the *z*_*c,g*_ := `logit`(*ψ*_*c,g*_) is a linear combination of ***x*** and ***y*** as follows,

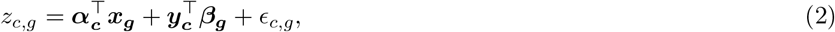

The model can be approximated deterministically by taking *ϵ*_*c,g*_ := 0, assuming all uncertainty comes from the regression weights. On the other hand, we could introduce 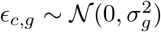 to account for gene-specific over-dispersion, which is particularly important for the potential phenomenon of mono-isoform in single cells.

Considering the overdispersion setting, we have a predictive distribution for *z*_*c,g*_, as follows

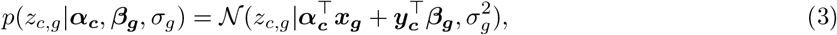

which can be treated as an informative prior on *z* (and accordingly giving rise to a logit-normal distribution for *ψ*).

### Bayesian Inference in BRIE2

Besides estimating the parameters for the regression model in Eq.(3), it is often of high interest to approximate the posterior distribution of the isoform abundance Ψ or its logit transformation *Z*. Therefore, it is crucial to keep *Z* as an auxiliary variable instead of marginalizing it out. By taking the product of the likelihood distribution defined in Eq.(1) and the predicted prior in Eq.(3), we could have the joint distribution to which the posterior distribution *p*(*Z*|*S*, A, B, ***σ***) is proportional as follows,

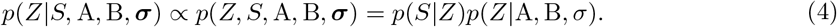

This posterior is intractable and it also has hype-parameters to optimize. In the BRIE1, we used an approximate algorithm to alternately optimize the parameters and sampling the posterior with Metropolis-Hastings algorithm [15]. Here, instead we are using a variational inference to approximate the posterior. Namely, we introduce a fully factorized distribution (mean-field) as a variational posterior, and we assume it is Gaussian, the same form as the predicted prior distribution in Eq.(3):

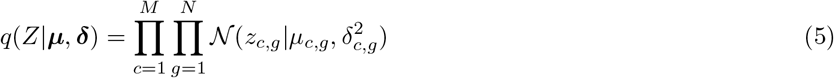

Then the inference becomes an optimisation problem for minimising the Kull-back–Leibler (KL) divergence between the exact Eq.(4) and variational posteriors Eq.(5),

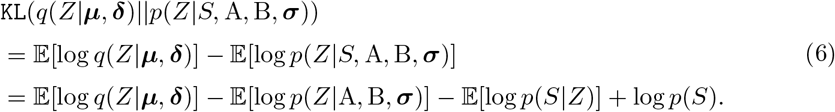

As the log *p*(*S*) is a constant term, minimizing the KL divergence is equivalent to maximizing the evidence lower bound (ELBO)

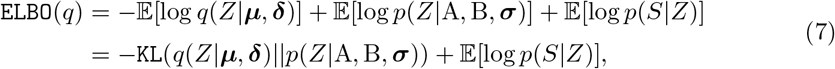

where 𝔼[·] denotes expectation over variational distribution *q*(*Z*) as a shortcut. The first part in ELBO is the `KL` divergence between the posterior and prior distribution on *Z*, which could be calculated analytically. The second term 𝔼[log *p*(*S*|*Z*)] in ELBO (Eq. (7)) is difficult to calculate due to the intrinsic mixture of two isoforms in the base likelihood Eq (1). Therefore, a cheap (but unbiased) Monte Carlo estimate [24] is introduced by sampling *R* samples of *Z* following its posterior distribution *q*(*Z*):

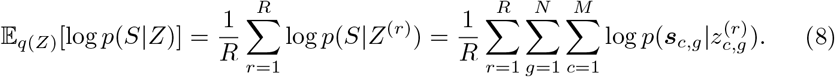

In practice, *R* = 3 samples are sufficient to give good estimates and are used by default. Given the expression of ELBO, we use a (stochastic) gradient descent algorithm, e.g., Adam by default [25], to achieve the maximum of ELBO. Here, we use the TensorFlow platform to obtain an automated derivation of the gradient. Also, the re-parametrization trick [24] for the gradient is fully supported for Gaussian distribution in TensorFlow.

### Detecting differential splicing events or differential momentum genes

BRIE2 (Mode 2-diff), in a unified way, allows to detection of differential alternative splicing events or differential momentum genes that are significant associated with one or multiple cell-level covariates. Notice that, since Mode 2 does not use sequence level features, different events will be independent under the model, therefore allowing model comparison on an event by event basis. This task is equivalently to select Model 1 (ℳ_1_) with significant effects (non-zero coefficient) versus Model 0 (ℳ_0_) with no effects (zero coefficient) for given cell feature(s) on a per event / gene basis. Therefore, BRIE2 will be run twice respectively for ℳ_1_ with all provided cell features and ℳ_0_ with leaving the candidate feature(s) out. For each event / gene *g*, we have the exact posteriors via the complete distributions 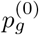 and 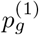, as follows,

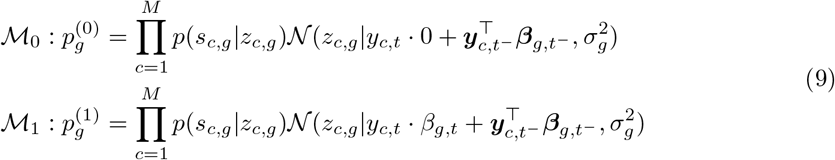

where *t*^−^ denotes all cell features except feature *t*. Then, we can run BRIE2 twice and obtain the optimised variational posteriors 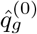 and 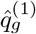, respectively. Then we compare the relevant evidence lower bounds `ELBO`_1_ and `ELBO`_0_, and obtain an `ELBO_gain` for event (or gene) *g* as follows,

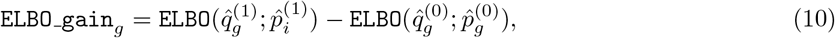

which approximates the empirical Bayes factor.

When testing events with alternative splicing associated with multiple sclerosis, we used the mouse strain, EAE state and intercept as covariates in ℳ_1_ and left EAE state out in ℳ_0_. When testing genes with splicing ratio associated to cell type, each time we include the proportion of detected genes, intercept, and one of 14 cell types as covariates in ℳ_1_ and left the cell type out in ℳ_0_. This test has been performed with OPC and OL as candidate cell type individually.

### Simulations

Simulations were performed to evaluate the quantification of *ψ* (Supplementary Fig. S5) and detection of splicing events that are significantly associated with cell-level features (main Fig. 2 and Supplementary Fig. S5-7). In both situations, we used an experimental data set with 130 cells and 2,248 splicing events [26] as seed data (see below).

Here, generate the *ψ* value with two steps. First, we quantified the *ψ* from the experimental seed data with 5 principal components as cell covariates. Then we calculated the mean *z* = `logit`(*ψ*) for each event across cells at 6.5 days, and their standard deviations (see Supp. Fig. S23). Then this observed PSI profile (averaged for one condition) will be used as a seed profile. We further generated the ***z*** profiles for the 130 cells through a Gaussian distribution by using the observed mean ***z*** and a fixed variance = 1, which well represents the experimental observations (Supp. Fig. S23). Second, we randomly but equally split the 130 cells into two conditions. We further randomly selected 400 out of 2,248 events as true differential alternative splicing (DAS) events, where we randomly plus or minus a fixed effect at size *η* to obtain an updated *z*′ = *z* ± *η* for the 65 cells in condition 2. We repeated with different effect size levels *η* at 2, 3, or 5 (see Fig. 2).

By using the same total read count of each event and cell as the seed data, we multiplied the simulated *ψ* to obtain the reads per kilobase (RPK) values for each isoform, which will be used as input for a sequencing read simulator Spanki [27]. Based on these generated reads, BRIE2 (with or without cell features), rMATS (v4.1.1), MAJIQ (v2.2) are performed to quantify PSI and detect the DAS between the two conditions.

### Data processing and gene filtering

For benchmarking BRIE2, we used 130 mouse embryonic cells at day 6.5 (80 cells) and day 7.75 (50 cells) that were generated [26] with SMART-seq2 protocol [28]. This data set has also been used as an illustration data set in BRIE1 [29]. Here, we used HISAT v2.2.0 [30] to align the reads to mouse genome GRCm38.p6, combined with ERCC92 spike RNAs. Then *brie-count* command line in BRIE v2.0.3 with all default parameters was used to count the reads aligned to each of the 8,253 alternative splicing events, which was extracted from GENCODE vM24 by using *briekit* at lenient thresholds.

The same processing except removing ERCC92 reference was applied to another SMART-seq2 data set on 2,208 mouse cells in the topic of multiple sclerosis [7], where BRIE2 was used to detect differential alternative splicing between disease and control cells. Here and in general, where detecting differential splicing and only cell-level features are applicable, we filtered out clearly less informative genes. By default in *brie-quant*, we filter out events with 1) less than 50 total reads or 10 unique reads across all cells, or 2) less than 30 cells with unique reads, or 3) the fraction of unique reads on minor isoform less than 0.001.

For RNA velocity analysis, a data set on dentate gyrus development was used, which was generated [31] with droplet protocol with 10x Genomics platform. The cell type annotation, UMAP visualization coordinates, and processed count matrices for both spliced and unspliced RNAs across 2,930 cells and 13,913 genes were downloaded from the tutorial in scVelo [9]. Only the top 3,000 highly variable genes with a minimum of 30 shared counts were used as suggested by scVelo. For detecting differential momentum genes, we only kept genes that were detected with at least one read in >15% of the cells. ScVelo v0.2.1 downloaded from PyPI is in use.

Additionally, an scNT-seq data set on excitatory neurons were obtained from the original paper [13]. This processed data set has 3,066 quality controlled cells and 44,021 genes. It also has UMAP visualization coordinates, time annotation and layers of spliced, unspliced, and new RNAs. Therefore, no pre-processing is needed on this data set. The RNA velocity inference by Dynamo (v0.95.2.dev142+9c30240) is based on the same scripts provided in the original paper [13]. When running the scVelo dynamical model, we select genes with at least 30 shared counts and either top 2,000 highly variable genes (Fig. 4d, Supplementary Fig. S17a and S18-S21) or top 8,000 highly variable genes though with only 4880 genes pass the requirement (Supplementary Fig. S17b).

### Quantitative metric for assessing RNA velocity

To evaluate the correctness of estimated cell transition from RNA velocity, we adopted the score of cross-boundary correctness proposed in [32]. Briefly, it measures the averaged cosine score between the velocity vector and delta expression vector of two groups of cells with known transition direction (from group A to B), as follows,

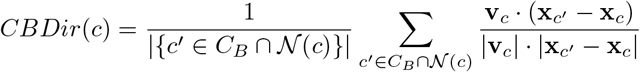

where cell *c* is from group A, 𝒩 (*c*) denotes the neighbour cells of *c*, and *c*′ ∈ *C*_*B*_ means a cell from group B. The **v**_*c*_ and **x**_*c*′_ − **x**_*c*_ respectively denote the velocity and delta expression vectors (usually in a reduced dimensionality space, UMAP is used here).

## Availability of Data and Materials

BRIE2 is an open-source Python package available at https://pypi.org/project/brie/ with Apache License 2.0. The version 2.0.5 used for analysis in the paper is deposited in Zenodo [33]. All the analysis notebooks and the processed data sets can be found at https://github.com/huangyh09/brie-tutorials. BRIE2’s manual with examples is available at https://brie.readthedocs.io.

## Authors’ contributions

Both authors conceived the study, carried out the data analysis, and wrote the paper. YH developed the software. Both authors read and approved the final manuscript.

## Ethics approval and consent to participate

Not applicable.

## Additional File

Additional file 1 — Supplementary Table S1 and Supplementary Figures S1–S22.

## Footnotes

[1] Notice also that using only unambiguous reads can significantly bias estimation for very low coverage levels, see Supp. Fig. S3 for the impact of using ambiguous reads on the PSI likelihood.

[2] Note, here we used the difference of estimated mean PSI in two conditions (i.e., delta_PSI) as indicators for both rMATS and MAJIQ, because the reported *p* values perform significantly worse, especially for rMATS (Supp. Fig. S7). This lower effectiveness of statistical significance may be due to the high sparsity of read counts in scRNA-seq data.

