## Supplementary File for "Computational identification of splicing phenotypes from single cell transcriptomic experiments"

### Supplementary materials for "Computational identification of splicing phenotypes from single cell transcriptomic experiments"

Yuanhua Huang and Guido Sanguinetti

#### 1 Supplementary Table

Table S1: The score of cross-boundary direction correctness (see Methods) between cell groups A to B. In the neural genesis data set (Dentate Gyrus, DG), the expected cell direction is from OPC to OC, and in the neural stimulation data set (scNT), the expected directions are 0 to 15min, 0 to 30min, 15 to 30min, 30 to 60min and 60 to 120min. For scNT data, an averaged score between multiple subgroups is used for the overall performance. Bold values mean the highest score in a certain group of cells. Four gene selection methods are compared here: scVelo's default gene selection, edgeR's differentially expressed genes, scVelo's differential kinetic genes (with different levels of FDR), and BRIE2's differential momentum genes (with different levels of ELBO gain, an approximate of Bayes factor).

| | DG: OPC->OL | scNT: overall | scNT: 0->15 | scNT: 0->30 | scNT: 15->30 | scNT: 30->60 | scNT: 60->120 |
|---|---|---|---|---|---|---|---|
| scvelo-default | -0.949 | 0.221 | 0.163 | -0.068 | 0.259 | **0.302** | **0.447** |
| edgeR-DEG: FDR<1% | -0.011 | 0.343 | 0.426 | 0.474 | 0.379 | 0.231 | 0.204 |
| diff-kinetics: FDR<1% | -0.990 | -0.247 | -0.672 | -0.575 | -0.319 | 0.083 | 0.250 |
| *diff-kinetics: FDR<5%* | -0.984 | 0.421 | **0.743** | **0.639** | 0.423 | 0.263 | 0.039 |
| diff-kinetics: FDR<10% | -0.969 | 0.343 | 0.426 | 0.474 | 0.379 | 0.231 | 0.204 |
| BRIE2-DMG: BF>3 | 0.645 | 0.463 | 0.738 | 0.629 | **0.416** | 0.275 | 0.257 |
| *BRIE2-DMG: BF>5* | **0.892** | **0.473** | 0.742 | 0.632 | 0.411 | 0.287 | 0.292 |
| BRIE2-DMG: BF>7 | 0.626 | 0.471 | 0.739 | 0.631 | 0.411 | 0.292 | 0.281 |

#### 2 Supplementary Figures

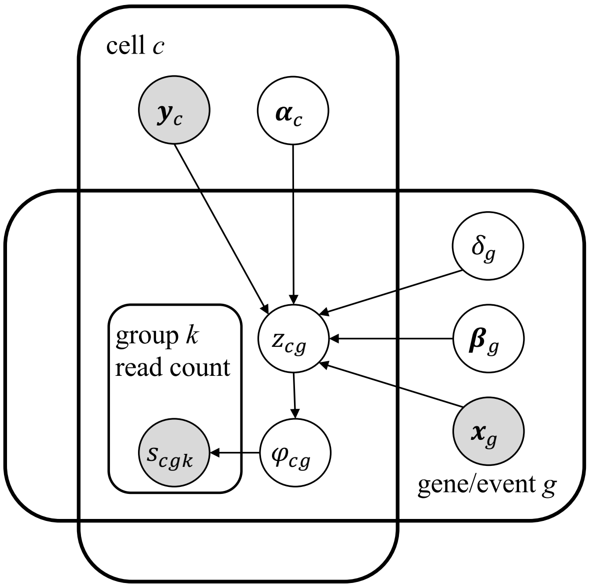


Figure S1: Graphical representation of BRIE2. In this model, the likelihood $p(\boldsymbol{s}_{c,g}|\psi_{c,g})$ of observing read count vector $\boldsymbol{s}_{c,g} = [s_{c,g,1}, s_{c,g,2}, s_{c,g,3}]$ is defined by a multinomial distribution whose parameter is based on length-weighted functions of $\psi_{c,g}$ (Methods). Here, as the main latent variable, $\psi_{c,g}$ denotes the proportion of isoform 1 (i.e, exon inclusion in skipping-exon event, or spliced RNA in velocity). It has a logit transformation to $z_{c,g}$ that has Gaussian prior with variance $\delta_g^2$ and mean $\boldsymbol{\alpha}_c^\top \boldsymbol{x}_g + \boldsymbol{y}_c^\top \boldsymbol{\beta}_g$ determinately by gene features $\boldsymbol{x}_g$, cell features $\boldsymbol{y}_c$, and their according coefficients $\boldsymbol{\alpha}_c$ and $\boldsymbol{\beta}_g$. Nodes in shade are observed, otherwise unknown variables.

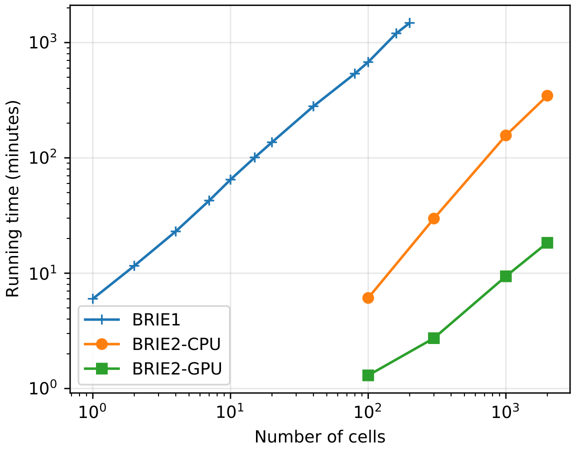


Figure S2: Running time of BRIE1 and BRIE2 (CPU or GPU) on 1 to 2,000 cells in a mouse multiple sclerosis data set (see main Fig. 3). BRIE1 runs on a Ubuntu server (with 80 CPUs at 2.00GHz and 376 GB memory), by using 25 CPU cores and it roughly consumes 8GB of memory on average. Given that it runs on cells separately, the running time is linearly according to the number of cells in the process. BRIE2-CPU runs on the same CPU server with a batch size of 30,000 elements, allowing CPU resources dynamically and automatically with Numpy and TensorFlow. It on average consumes 10-25 CPUs and around 4 GB of memory. BRIE2-GPU runs on another Ubuntu server with 4 Nvidia GPUs (GeForce RTX 2080 Ti), each with 11 GB of memory. It uses one GPU card and on average consumes 8GB of memory with the same batch size as on CPU server.

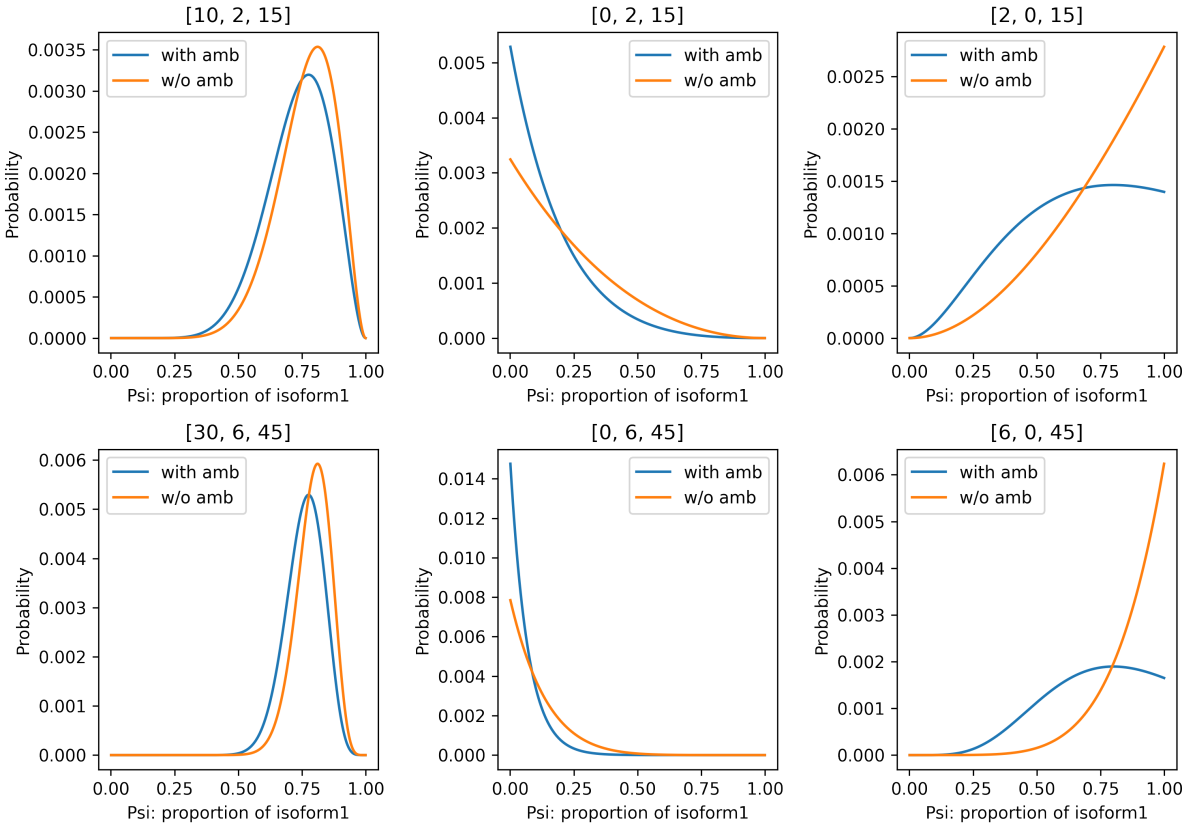


Figure S3: Examples of likelihood distribution by including (in blue) or excluding (in orange) ambiguous reads. The vector in the title of each subplot means isoform1-specific, isoform2-specific and ambiguous reads, receptively. Here the effective lengths of isoform1, isoform2 and ambiguous positions of this toy gene are 200nt, 100nt, and 500nt, respectively.

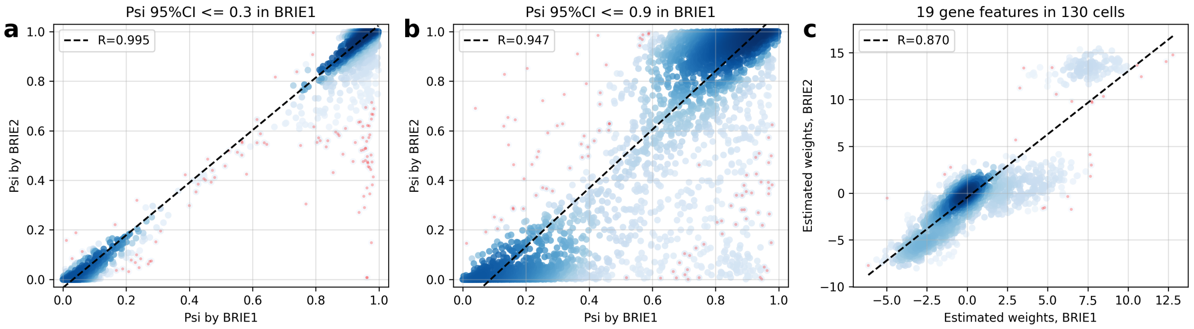


Figure S4: Comparison between BRIE1 and BRIE2 on 130 mouse embryonic cells. (a-b) Scatter plot of estimated PSI by BRIE1 and BRIE2 for those with 95% confident interval in BRIE1 $<0.3$ (a) or $<0.9$ (b). (c) Scatter plot of estimated weights on 19 gene features in each of the 130 cells between BRIE1 and BRIE2. R: Pearson's correlation coefficient.

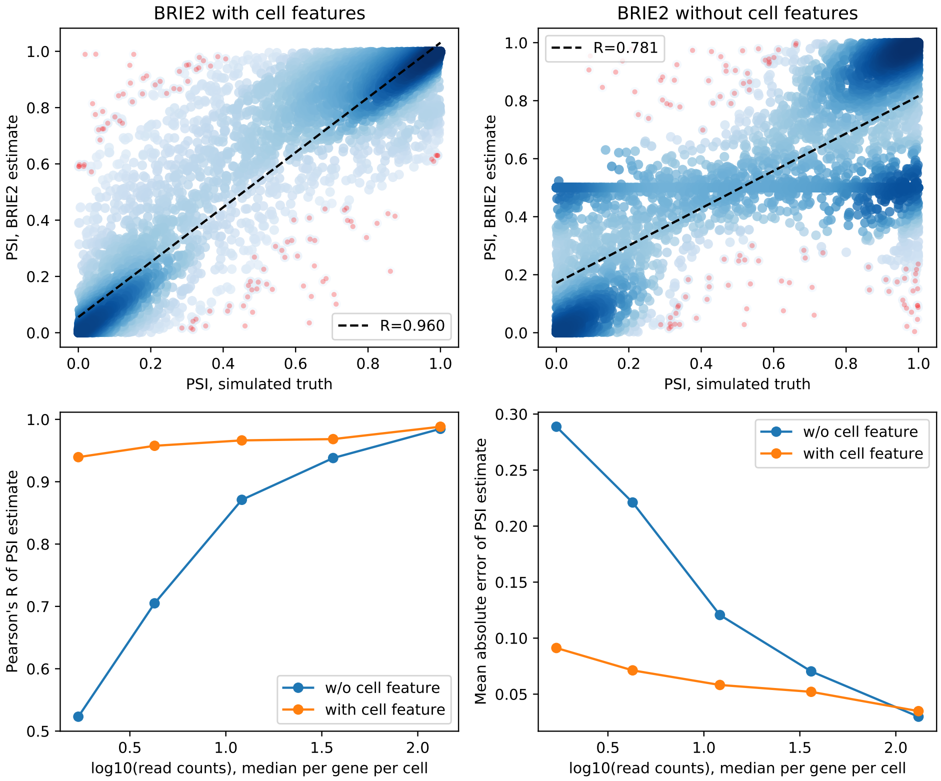


Figure S5: Cell feature based prior improves PSI quantification. (a-b) Comparison between BRIE1 and BRIE2 on 130 mouse embryonic cells. (a-b) Scatter plot of estimated and simulated PSI by including a unity cell feature for aggregated prior (a) or no feature at all (b). (c) Pearson's correlation coefficients between estimated and simulated PSI, at different expression levels. (d) Mean absolute error between estimated and simulated PSI, at different expression levels.

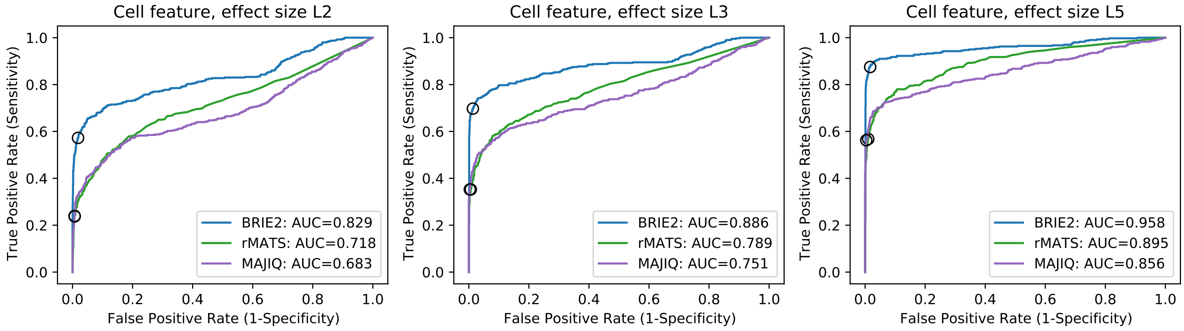


Figure S6: Receiver operating characteristic curves of detecting differential alternative splicing with different levels of effect size. This figure has the same settings of main Fig. 2.

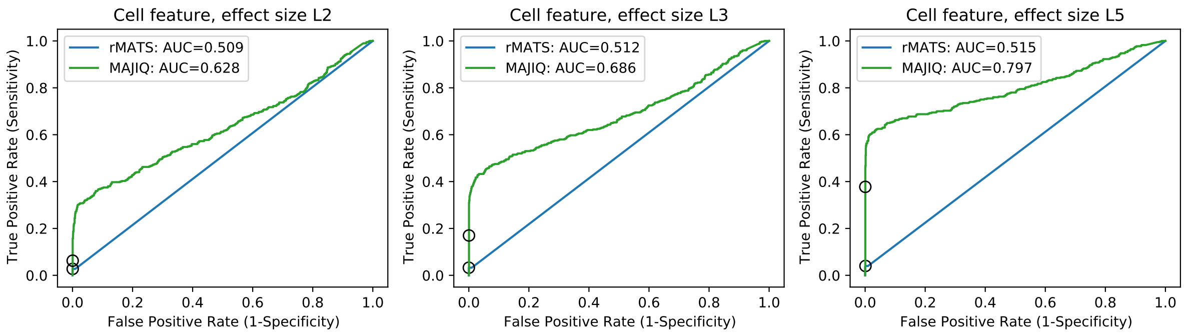


Figure S7: Receiver operating characteristic curves of detecting differential alternative splicing with different levels of effect size. Opposed to the use of |`delta_Psi`| in main Fig. 2 and Supp. Fig. S6, p values are used here as indicator for both rMATS and MAJIQ.

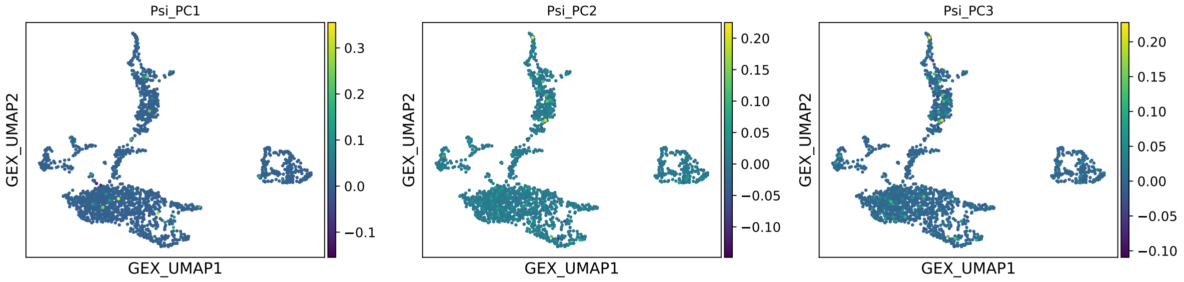


Figure S8: The first three principal components of PSI matrix on simulated data with homogeneous PSI vector for all cells and observed expression counts from real data. Here, PSI is quantified from the simulated reads by using a unity cell feature for aggregation (i.e., Mode 2-quant).

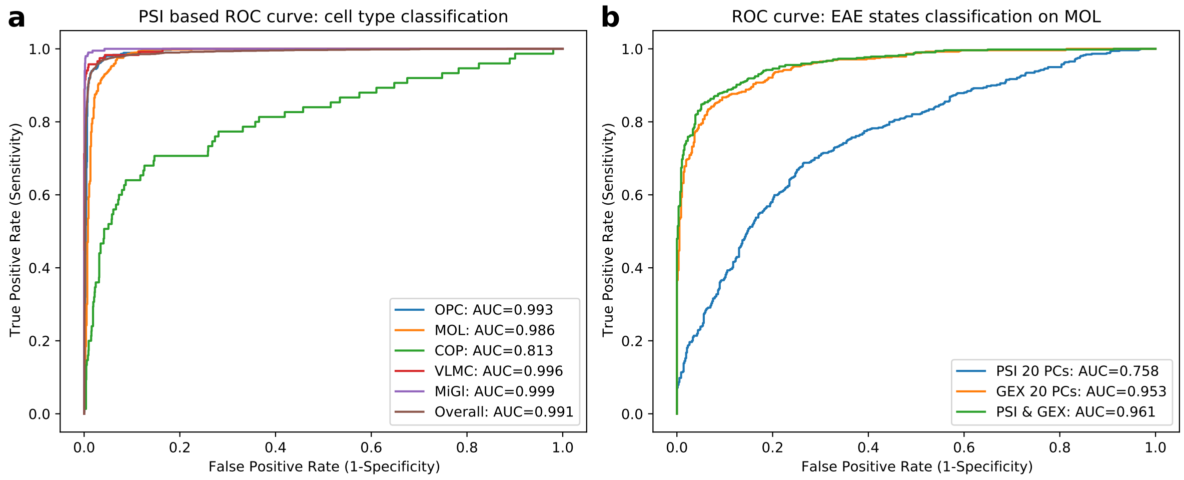


Figure S9: Prediction performance with PSI based principal components (PC). (a) Receiver operating characteristic curve for prediction of cell types from the first 20 PCs of PSI matrix by using logistic regression in a multi-label classification. Ten-fold cross-validation is used for the evaluation, where overall means combining all cell types in a micro-manner. (b) Receiver operating characteristic curve for prediction of EAE condition in mature oligodendrocytes (MOL) using logistic regression. Predictive features are: 1) the top 20 PCs of splicing proportion (i.e., PSI), 2) gene expression (i.e., GEX) or 3) both. Ten-fold cross-validation is used for the evaluation.

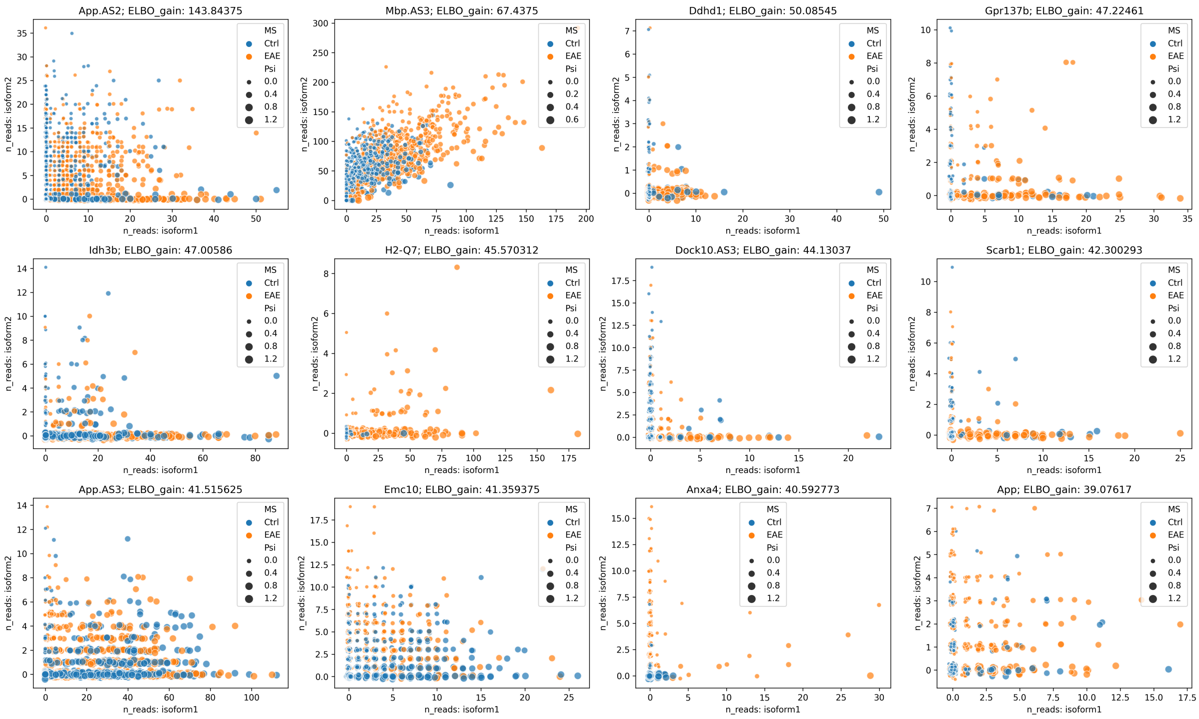


Figure S10: Scatter plots of read counts from isoform1 (x-axis) and isoform2 (y-axis) unambiguously on the 12 most significantly differential splicing events between EAE and control. Each subplot represents a gene with each dot referring to a cell.

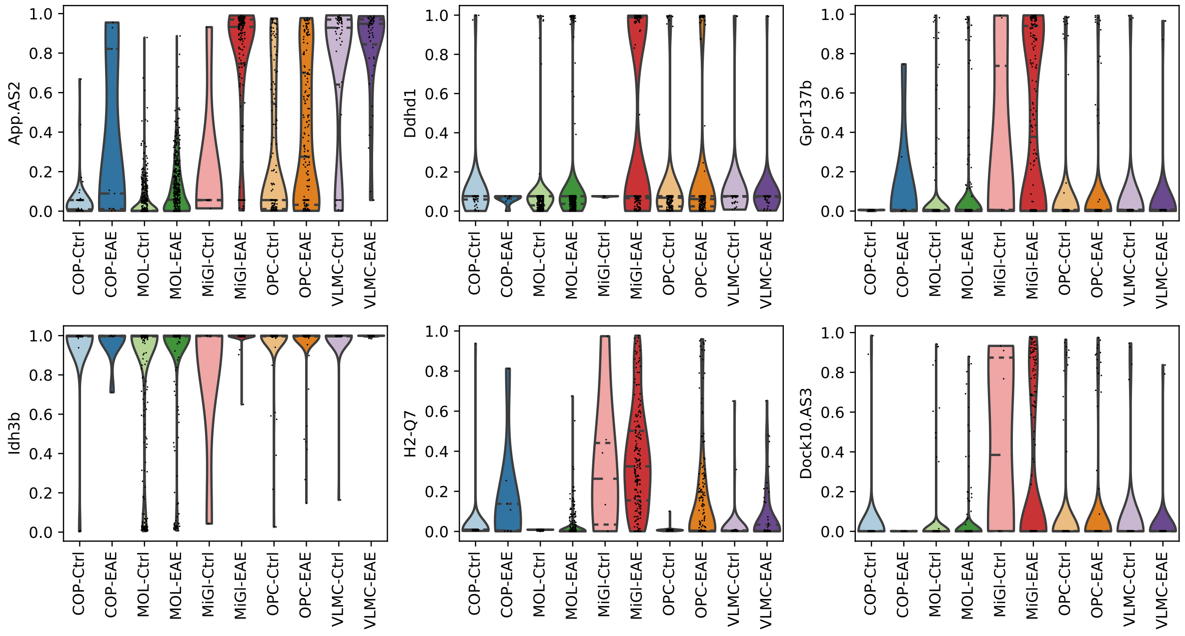


Figure S11: Violin plots of estimated PSI on 6 most significant splicing events between EAE and control. Each dot is a cell, which is categorised into EAE and control in each cell type. PSI values are quantified by only using unity cell feature for aggregation (i.e., Mode 2-quant).

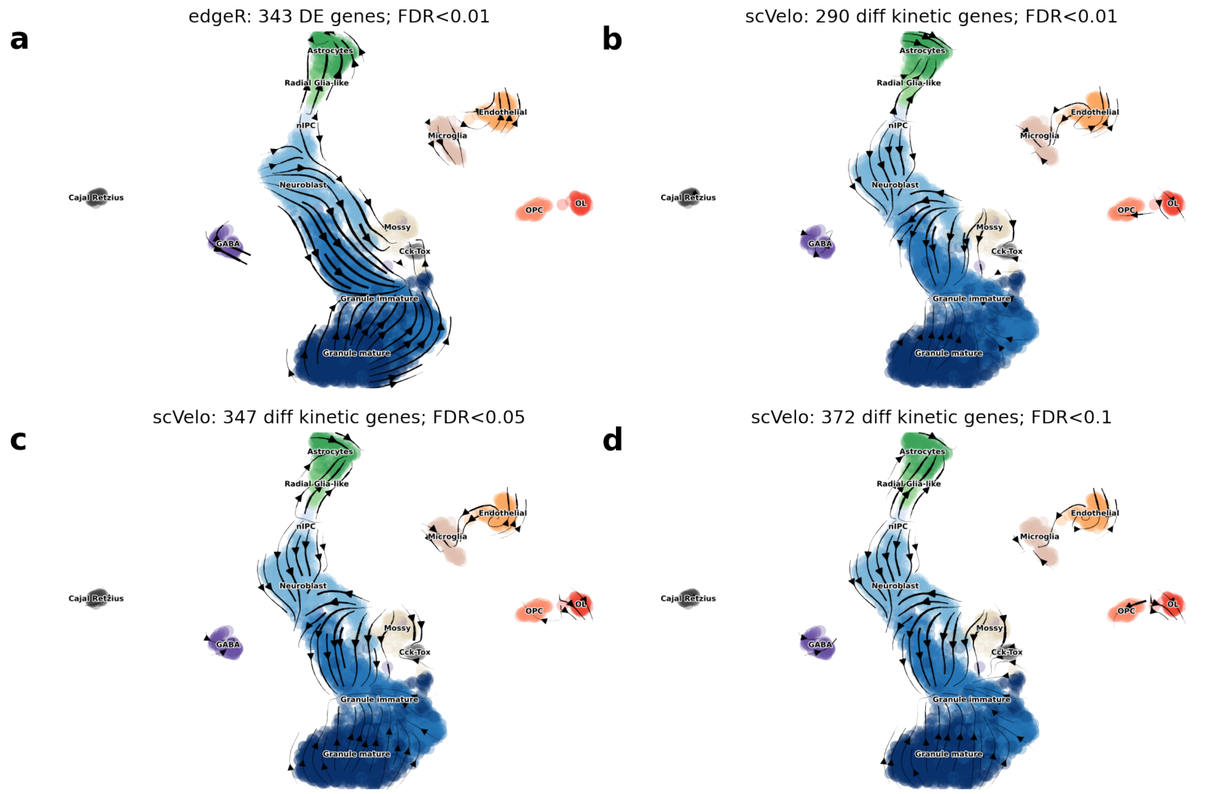


Figure S12: Cell differentiation in neurogenesis (Dentate Gyrus) data set inferred from RNA velocity by scVelo with different gene sets: (a) 343 differentially expressed genes by edgeR (FDR$< 0.01$); (b-d) genes with differential kinetic rates by scVelo at FDR$< 0.01$ for 290 genes (a), at FDR$< 0.05$ for 347 genes (b), and at FDR$< 0.1$ for 372 genes (d). All differential tests in (a-d) are performed on OL vs rest and OPC vs rest. This figure is related to main Fig. 4a.

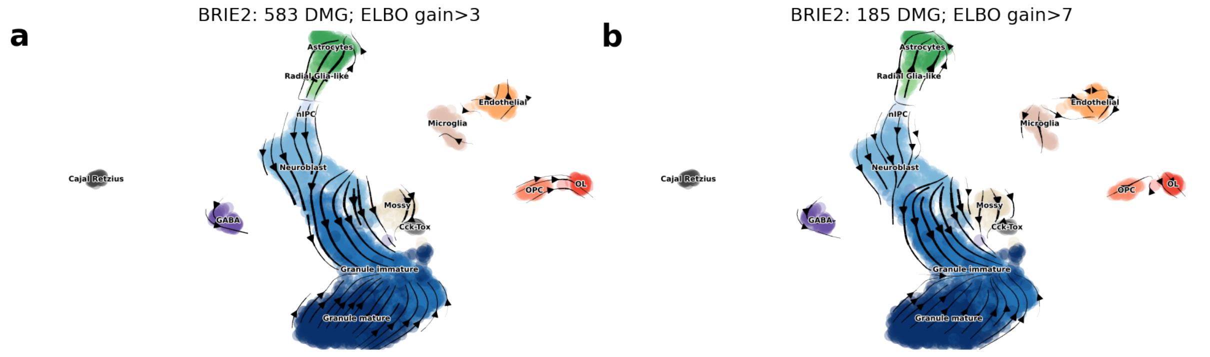


Figure S13: Cell differentiation in neurogenesis (Dentate Gyrus) data set inferred from RNA velocity by scVelo with different gene sets: (a) 583 differential momentum genes detected by BRIE2 (ELBO_gain>3); or (b) 185 DMGs (ELBO_gain>7). All differential tests in (a-b) are performed on OL vs rest and OPC vs rest. This figure is related to main Fig. 4b.

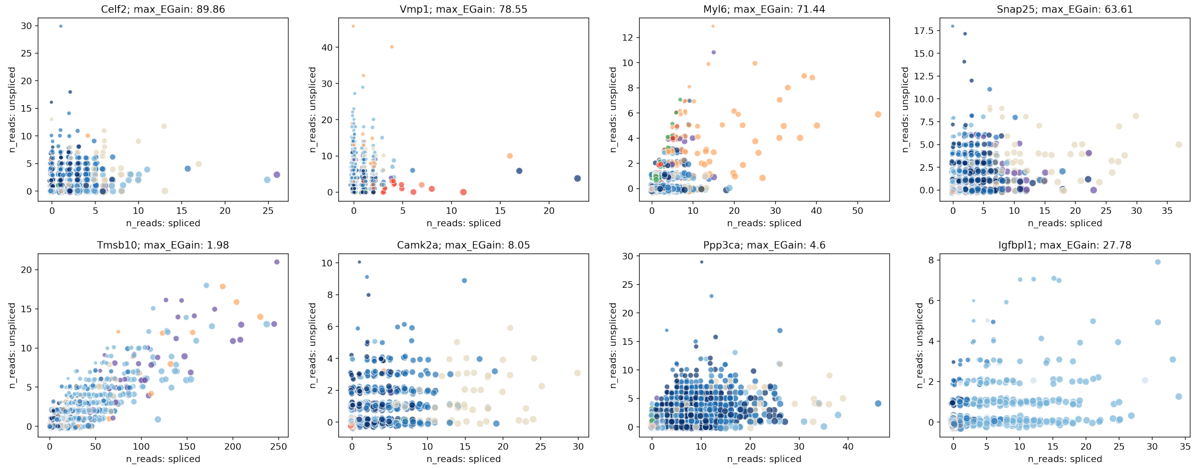


Figure S14: Scatter plots of read counts from spliced (x-axis) and unspliced (y-axis) RNAs on top four differential spliced genes between one cell type versus others detected by BRIE2 (top panel), and four genes that were picked in scVelo paper (bottom panel).

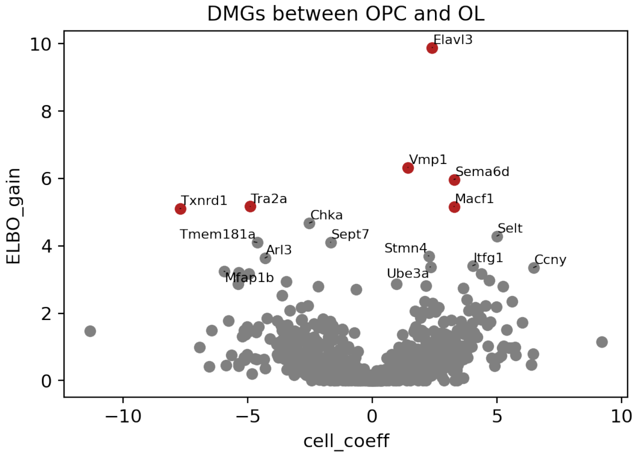


Figure S15: Volcano plot of BRIE2 detected differential momentum genes between oligodendrocyte precursor cells (OPC) and myelinating oligodendrocytes (OL). Shown is ELBO_gain on the y-axis and the effect size of logit(Psi) on the x-axis. PSI means the proportion of spliced RNAs and positive effect size means a higher proportion of spliced RNAs in OL. Note, only the 103 OPC and OL cells are used for this analysis, instead of full dataset in Supp. Fig. S13.

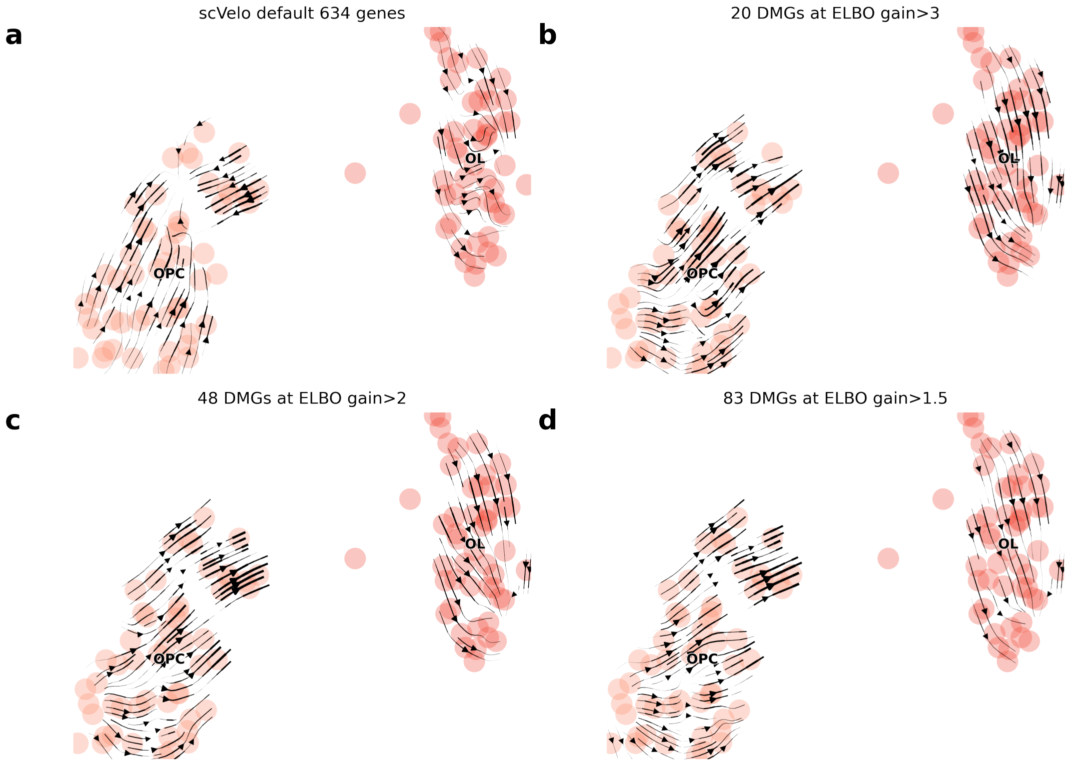


Figure S16: Cellular transition among 103 OPC and OL cells inferred from scVelo with different gene sets: (a) 634 gene detected by scVelo; (b) 20 differential momentum genes (DMGs) detected by BRIE2 with ELBO_gain $> 3$; (c) 48 DMGs with ELBO_gain $> 2$; (d) 83 DMGs with ELBO_gain $> 1.5$. The expected direction is from OPC to OL, which is consistent with DMGs across different thresholds.

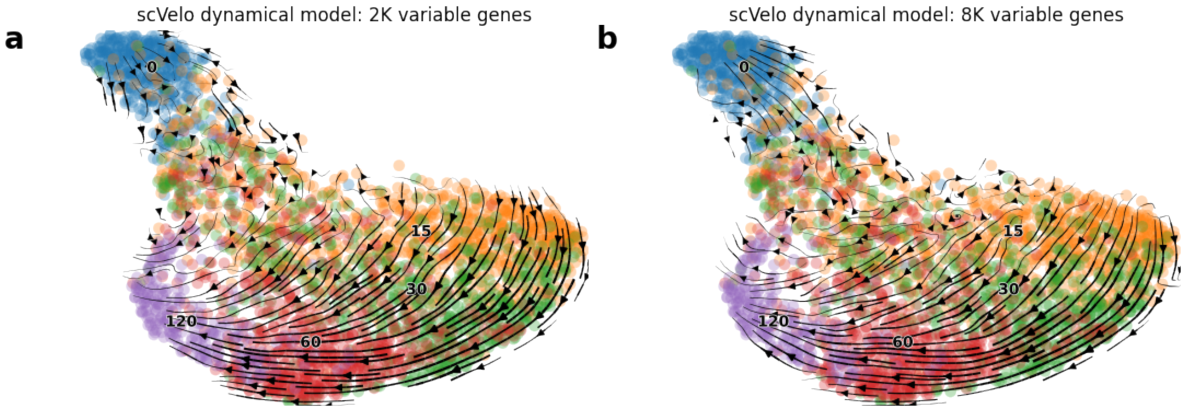


Figure S17: Cell transitions of excitatory neurons inferred from RNA velocity on total RNAs by scVelo with a different initial number of highly variable genes. (a) Top 2,000 highly variable genes, resulting in 131 genes with effective RNA velocities or (b) Top 8,000 highly variable genes (only 4880 passing minimum shared 30 counts), resulting in 251 genes with effective RNA velocities for projecting cellular transitions.

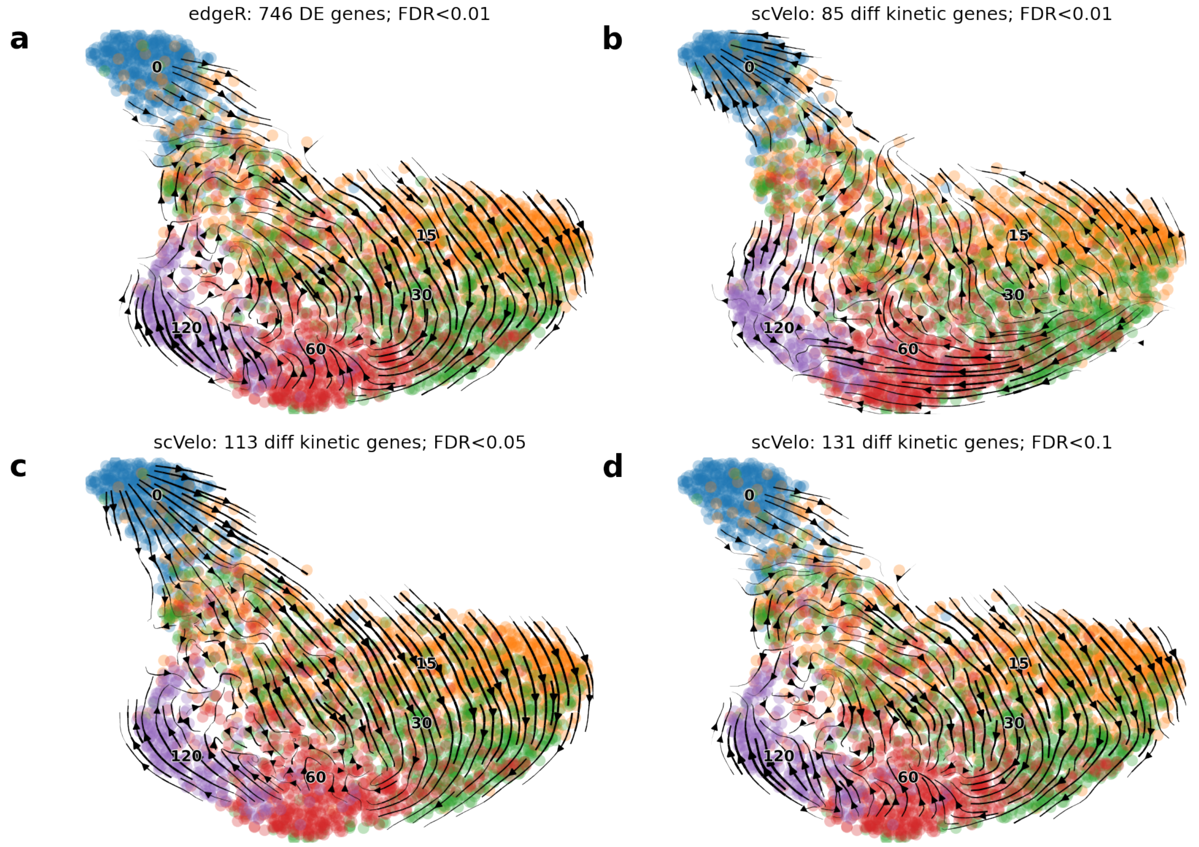


Figure S18: Cell transitions of excitatory neurons inferred from RNA velocity on total RNAs by scVelo with 2000 initial number of highly variable genes and further filtered by using (a) 746 genes with differential expression along time by edgeR (FDR$< 0.01$); (b-d) genes with differential kinetic rates at any time point by scVelo at FDR$< 0.01$ for 85 genes (b), at FDR$< 0.05$ for 113 genes (c) and at FDR$< 0.1$ for 131 genes. This figure is related to main Fig. 4d.

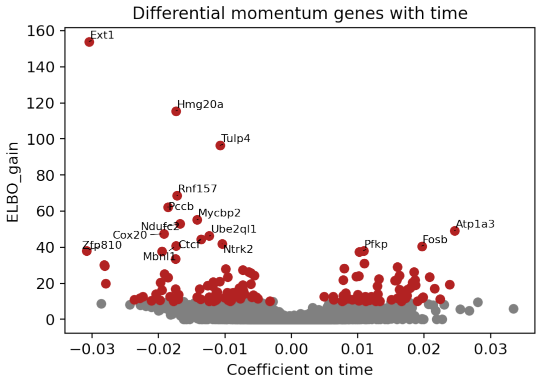


Figure S19: Volcano plot of BRIE2's detected differential momentum genes associated with stimulation time on excitatory neurons. Shown is ELBO_gain on the y-axis and effect size of logit(Psi) on the x-axis for detecting differential splicing ratio between stimulation time ranging from 0 to 120 minutes.

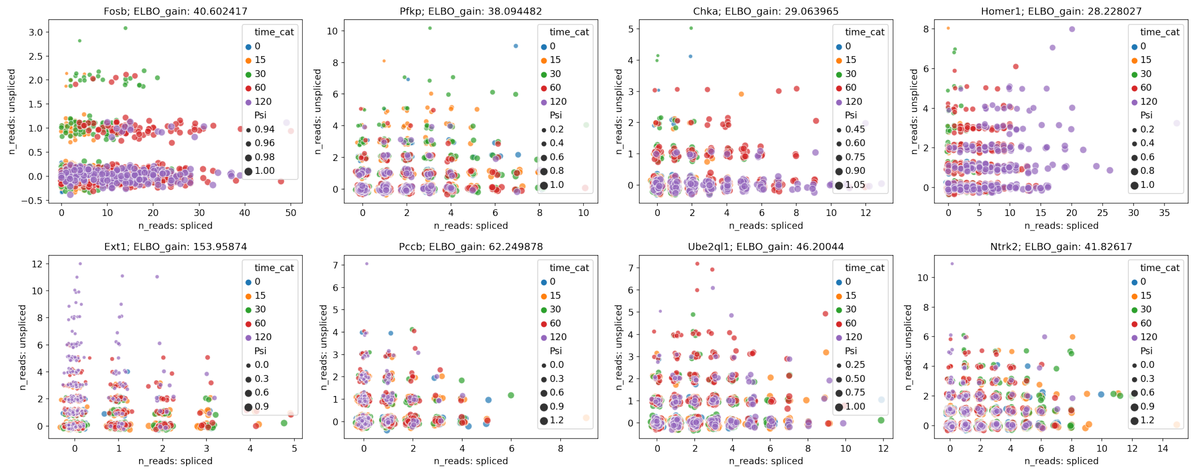


Figure S20: Scatter plots of read counts from spliced (x-axis) and unspliced (y-axis) RNAs on eight differential momentum genes detected by BRIE2, namely splicing ratio significantly associated with stimulation time. Top panel for top four positively affected genes and bottom panel for top four negatively affected genes.

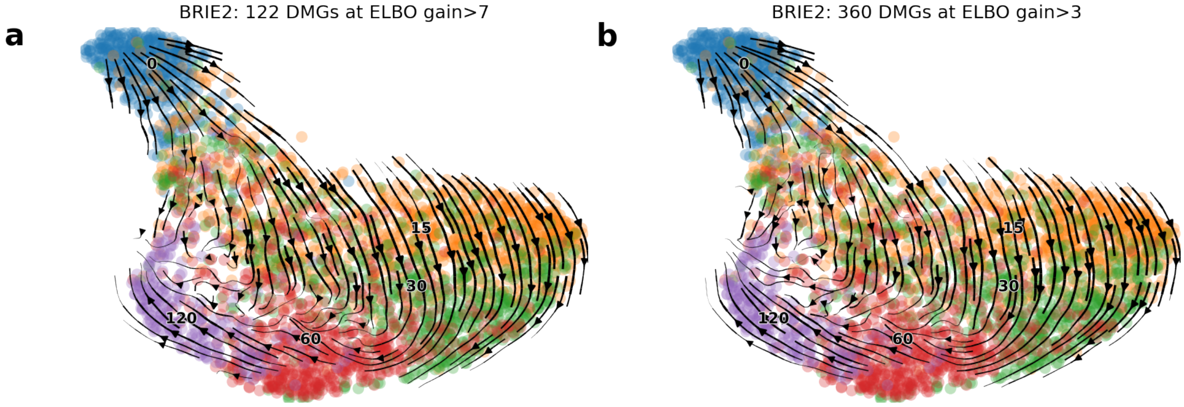


Figure S21: Cell transitions of excitatory neurons inferred from RNA velocity on total RNAs by scVelo with (a) 122 differential momentum genes detected by BRIE2 (ELBO_gain>7); or (b) 360 DMGs (ELBO_gain>3);

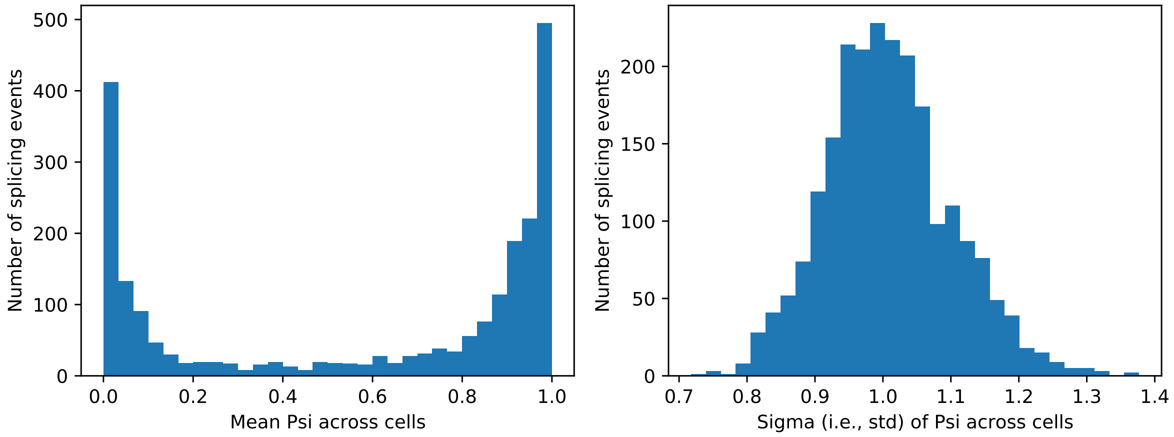


Figure S22: Distributions of key parameters of 2,248 splicing events that are observed from experimental seed data for simulation: (a) mean PSI and (b) standard deviation of PSI both across 80 cells at day 6.5. Simulation results are in main Fig. 2 and Supp. Fig. S5-7.
